# How important is budding speciation for comparative studies?

**DOI:** 10.1101/2022.05.24.493296

**Authors:** Daniel S. Caetano, Tiago Bosisio Quental

## Abstract

The acknowledgment of evolutionary dependence among species has fundamentally changed how we ask biological questions. Phylogenetic models became the standard approach for studies with three or more lineages, in particular those using extant species. Most phylogenetic comparative methods (PCMs) translate relatedness into covariance, meaning that evolutionary changes before lineages split should be interpreted together whereas after the split lineages are expected to change independently. This clever realization has shaped decades of research. Here we discuss one element of the comparative method often ignored or assumed as unimportant: if nodes of a phylogeny represent the dissolution of the ancestral lineage into two new ones or if the ancestral lineage can survive speciation events (i.e., budding). Budding speciation is often reported in paleontological studies, due to the nature of the evidence for budding in the fossil record, but it is surprisingly absent in comparative methods. Here we show that many PCMs assume that divergence happens as a symmetric split, even if these methods don’t explicitly mention this assumption. We discuss the properties of trait evolution models for continuous and discrete traits and their adequacy under a scenario of budding speciation. We discuss the effects of budding speciation under a series of plausible evolutionary scenarios and show when and how these can influence our estimates. We also propose that long-lived lineages that have survived through a series of budding speciation events and given birth to multiple new lineages can produce evolutionary patterns that challenge our intuition about the most parsimonious history of trait changes in a clade. We hope our discussion can help bridge comparative approaches in paleontology and neontology as well as foster awareness about the assumptions we make when we use phylogenetic trees.

---

Phylogenetic trees are the main representation of evolutionary relationships among lineages and stand as a symbol of evolutionary thought. However, their relatively simple structure is not capable of informing us about all aspects of evolution. Processes such as horizontal gene transfer, hybridization, and introgression can produce complex evolutionary relationships that challenge the explanatory power of bifurcating phylogenies (Philippe and Douady, 2003; Mallet et al., 2016; Bastide et al., 2018). Interestingly, the graphical representation of bifurcating trees could even influence how we think (Baum et al., 2005). A node connecting one ancestral branch to two new ones can suggest evolutionary histories much simpler than what we observe in nature and potentially downplay important aspects of macroevolution. For instance, if read literally, bifurcating phylogenies can be seen as speciation events that happened due to the split of an ancestral lineage into two new ones, coincident with the extinction, or dissolution, of the ancestral lineage (Meier and Willmann, 2000; Bokma, 2008). This view makes the concept of a lineage synonymous with a branch of a phylogenetic tree. However, evidence from empirical systems shows the mode of speciation can be varied and often complex (Rosenblum et al., 2012), including frequent instances of budding speciation (Wagner, 1998; Funk and Omland, 2003; Gottlieb, 2004; Crawford, 2010; Anacker and Strauss, 2014; Otero et al., 2019; Patsis et al., 2021).

The paleontological literature often adopts the representation of budding trees (e.g., Raup and Gould, 1974; Raup, 1985; Foote, 1996; Wagner, 1998; Benton and Pearson, 2001) that inform which branches are new lineages and which are the continuation of the ancestral lineage. Budding is recognized in the fossil record as a cladogenetic event in which a new lineage appears as a branching of an older lineage that can still be found after the speciation event (e.g., Foote, 1996). Although budding speciation is commonly reported in paleontology, it is rarely incorporated into phylogenetic comparative models (PCMs). Bokma and colleagues have developed a series of PCMs incorporating budding and the effect of punctuated equilibrium (Bokma 2002; Bokma, 2008; Matilla and Bokma, 2008; Monroe and Bokma, 2009; Bokma, 2010; Jansen et al., 2022). Unfortunately, these and other similar methods (Bartoszek, 2014; Bartoszek, 2020, Pagel et al., 2022) have not been widely used in the PCM literature, perhaps due to the perception that bifurcating molecular phylogenies show no evidence of budding speciation. Here we bring a different, and perhaps controversial, point of view; that budding speciation can be common, that it might affect inferences of trait evolution, that it can be detected in molecular phylogenies, and it should be considered in PCMs even when there is no information from the fossil record evidencing its role in the diversification of the group.

There has not been a consensus about the effect of budding speciation on our estimates of the tempo and mode of trait evolution using PCMs. Bokma (2008) implemented a trait evolution model to estimate the contribution from saltational and anagenetic changes, however, there is no investigation of the impact of the saltational process in inferences using PCMs that do not accommodate such effects. De Lisle and colleagues (2021) explored the effect of shifts in the adaptive optima on extinction rates using a model based on population-level dynamics and showed that lineages would rarely survive peak shifts and those occupying stable optima are expected to have a higher chance of survival. In turn, Duchen and colleagues (2021) studied how cladogenesis changes the average species phenotype using individual-level simulations and showed that new lineages budding off from an ancestral population can show a significant phenotypic deviation due to neutral processes. Combined, these results point to the idea that long-lived evolutionary lineages might occupy stable optima (Eldredge and Gould, 1972; Gould and Eldredge, 1993; also see Goldberg and Foo, 2020) whereas new lineages may bud off with distinct phenotypes, due to the effect of cladogenesis on the species trait and selection to occupy a distinct region of the morphospace (Eldredge and Gould, 1972; Gould and Eldredge, 1993; Bokma, 2008; Matilla and Bokma, 2008).

Here we discuss in which circumstances budding speciation can affect our estimates of the tempo and mode of trait evolution. More specifically, we review the properties of PCMs of trait evolution that dictate whether or not budding speciation can influence our conclusions. In our view, most PCMs of trait evolution were developed with the strong assumption that speciation is symmetric, lineages cannot continue after speciation events, and there is no effect of budding in molecular phylogenies. Here we discuss how budding speciation can bias our estimates using simulations as an argumentative guide to our narrative. Distinct from previous studies (Matilla and Bokma, 2008; De Lisle et al., 2021; Duchen et al., 2021; Crouch et al., 2021 among others), our discussion focuses on cases in which budding speciation is not considered when using PCMs.

## What is budding and how to recognize it?

Budding is defined as a speciation event in which the new species co-occurs in time with its ancestral lineage (Foote, 1996), meaning the ancestral lineage continues to exist after speciation. We use the term progenitor lineage to help differentiate the parent from the daughter lineage (see Gottlieb, 2004). A progenitor lineage is a lineage that has given birth to one or more new lineages through budding. Budding speciation is inherent in Mayr’s (1942) concept of speciation via peripatric speciation. Also, as discussed by Eldredge and Gould (1972) and Grant (1981), drift likely has an important impact on small founder populations and, naturally, will have consequences for trait evolution (see De Lisle et al., 2021). Perhaps due to its intrinsic role in diversification, the neontological literature has attributed different nomenclature to what is fundamentally budding speciation. In this section, we discuss how budding speciation has been recognized in the literature, which patterns might be the result of past budding events, and how budding can be detected using data from extant species. The reader will note that we attribute a variety of processes to the effect of budding speciation. Indeed, one of the main goals of this discussion is to bring awareness to the role of budding speciation in studies of macroevolution and how it connects to multiple patterns we observe in phylogenetic trees.

In the absence of the fossil record, budding has been recognized as a new lineage formed within or at the edge of the ancestral lineage (Anacker and Strauss, 2014) or as a biological cause of paraphyletic species (Funk and Omland, 2003; also see Fig. S1). This has also been associated with the hypothesis of Punctuated Equilibrium (Eldredge and Gould, 1972; Gould and Eldredge, 1993), since peripheral populations can become isolated and speciate, leading to fast trait evolution due to a shift in the optima for the traits (Mayr 1942; Eldredge and Gould, 1972; Grant, 1981; Gould and Eldredge, 1993; Bokma, 2008; Matilla and Bokma, 2008; De Lisle et al., 2021). Budding speciation is considered to be opposed by bifurcation—the split of an ancestral lineage into two new ones (Fig. S1). Hagen and colleagues (2015), for example, utilized the term *symmetrical* speciation to capture the role of allopatric speciation and opposed it to *asymmetrical* speciation which represents peripatric speciation with the continuation of the progenitor species—thus, budding. Although these authors use distinct nomenclature, each is an example of budding. The paleontological literature suggests budding is a common evolutionary pattern and some argue it represents the majority of speciation events observed in deep time. For example, Wagner (1998) used budding speciation to estimate a phylogeny of hyenas that provided a better fit to the stratigraphic history of the fossil record. Aze and colleagues (2011) reconstructed a large phylogeny of macroperforate foraminifera in which most cladogenetic events were recognized as budding through analysis of morphological characters. Bapst and Hopkins (2017) applied an explicit probabilistic model to date a phylogeny of trilobites and also show that budding events are often supported by the fossil record. Similarly, Parins-Fukuchi (2021) re-evaluated the diversification of hominins and suggests the occurrence of budding speciation events. In contrast, there is little to no mention of budding speciation in the neontological literature which, in our view, creates an undesirable disconnect between paleontology and neontology. One could argue that this absence is due to the impossibility of detecting budding using molecular phylogenies, however, as we discuss below, we disagree with this statement.

If budding speciation is frequent, we expect to recover recent events of budding using molecular data. When a new lineage buds off from its ancestral, the progenitor species becomes paraphyletic (Funk and Omland, 2003; see Fig. S1B). The advantage of neontological data is that molecular phylogenies can show evidence of budding independent of the use of morphological divergence to estimate the tree, which is necessary to both estimate phylogenies and detect speciation based on fossil remains (Foote, 1996; Wagner, 1998; Bapst, 2013). If the new lineage maintains cohesion and does not go extinct (see De Lisle et al., 2021) or is not reabsorbed via hybridization with the progenitor (Taylor et al., 2006; Richmond and Jockush, 2007; Behm et al., 2010; Lackey and Boughman, 2017), gene flow among the populations of the progenitor should complete sorting and the progenitor and daughter lineages will eventually become sister species in estimated molecular phylogenies—erasing the signal of budding. Thus, budding speciation can be detected using molecular phylogenies, but its signal disappears over time whereas, in the fossil record, the information is preserved if the record is reasonably complete. Otero and colleagues (2019) show an interesting case in *Iberodes* plants which underwent two events of budding within the last 5 million years. In both instances, the new lineage evolved distinct morphological and ecological traits (Otero et al., 2019). *Iberodes* has inland and coastal species and the potential change in selective pressure together with the peripheral distribution of the younger coastal lineages likely were key factors for budding divergence. Similarly, Papuga and colleagues (2018) show peripheral plant populations that have lower niche breadth (i.e., are more specialized) than central populations as well as divergence in ecological traits (i.e., soil parameters), both factors that can cause budding by ecological speciation. Strong evidence for budding speciation was also detected from molecular phylogenies by Baldwin (2005), showing that *Layia glandulosa* (a Compositae plant) is the progenitor species for *L. discoidea*. Anacker and Strauss (2014) tested 71 sister pairs and demonstrated that young divergences frequently show overlapping and asymmetrical ranges— another indication of budding speciation. This asymmetry was not detected among older clades, suggesting the signal of budding on the geographic distribution of sister pairs is lost as lineages get older. We also note that taxonomic revisions that re-name paraphyletic species into several new species, also erase the signal of budding.

If budding is frequent, and we suspect it is, it can be an important factor in understanding trait evolution because peripheral populations can show distinct mean phenotypic values (Papuga et al., 2018) and divergence through budding can generate new lineages with distinct average phenotypes (Gottlieb, 2004; Duchen et al., 2021) and evolutionary trajectories (De Lisle et al., 2021). If we assume molecular phylogenies are literal bifurcating trees, despite the evidence for budding speciation, then PCMs might be based on inadequate assumptions. In the next two sections, we visit the most popular PCMs and discuss scenarios in which the presence of budding would, or would not, affect our estimates.

## When budding doesn’t matter

Raup (1985) stated that budding should not influence estimates of net diversification rate because the addition or subtraction of lineages at any given time would be perceived similarly if we represent a phylogeny either by budding or bifurcation. This question has been revisited by Bapst and Hopkins (2017) and Crouch et al. (2021), both showing that budding can change divergence time estimation and alter estimates of the accumulation of lineages through time (also see Wagner, 1998). Thus, budding should not influence the net diversification rate only if the true dated tree is known, otherwise, changes in divergence time estimation can potentially impact estimates of diversification down the line.

With respect to models of trait evolution, budding should not influence our estimates if changes happening at any point in time, and at any branch of the phylogeny, are independent of the prior history of the lineage and their ancestors. Two important simplifications were introduced when Felsenstein described the method of independent contrasts (1981; 1985); trait changes happen independently in each branch of the phylogeny and evolutionary changes at each point along a branch are independent and identically distributed (iid). Most models of trait evolution share these assumptions (see review in O’Meara, 2012; Pennell and Harmon, 2013). However, most, if not all, PCMs were not created with the intent to accurately describe evolution in a mechanistic way, and the use of simplifications does not mean we assume evolution follows these rules.

If models of trait evolution that assume a homogeneous process across all branches of the tree are adequate representations of macroevolution, the incorporation of budding speciation will not change our estimates. This is because differentiating lineages in the phylogenetic tree will have no influence on the underlying model—lineages become effectively interchangeable. However, this is not the trend that we are currently observing in PCMs. Extensions allowing heterogeneity in the process, often associated with some predictor, have been shown to better capture the variation of empirical data (e.g., Eastman, 2011; Rabosky et al., 2014; Uyeda and Harmon, 2014; Caetano et al., 2018; Pagel et al., 2022). More recently, studies have demonstrated that rate heterogeneity should be taken into account even when no a priori predictors are present (e.g., Rabosky and Goldberg, 2015; Beaulieu and O’Meara, 2016; Caetano et al., 2018; May and Moore, 2020). Development of more adequate models often means the increase in model complexity to reflect the dynamic nature of macroevolution and, as a result, hint that the condition of homogeneous and memoryless evolutionary changes with interchangeable lineages—under which budding would not matter—is unlikely across the tree of life. Below we discuss how budding could be generating heterogeneity in the phylogenetic history of phenotypes and in which ways the results affect our conclusions about trait evolution.

## When budding matters

Budding is expected to be important in any evolutionary scenario in which the identity of evolutionary lineages is relevant. This might be the case if lineage age influences the tempo and/or mode of trait evolution (Hagen et al., 2018; Goldberg and Foo, 2020) or if the age of competing lineages is important to predict their competitive strength and/or risk of extinction (Ezard et al., 2011; Rosenblum et al., 2012; Carrillo et al., 2020; Januario and Quental, 2021). Although there are other evolutionary processes under which the identity of lineages might be important, here we focus on these two scenarios for simplification. In contrast, there are special cases that generate heterogeneity in trait evolution but under which budding likely is not relevant. For example, if shifts in rates of trait evolution are due to abiotic causes equally affecting all lineages concurrent with the event, such as response to climatic changes or mass extinctions, then, everything else being equal, we would expect responses to be independent of lineage identity (e.g., Clavel and Morlon, 2017). Below we enumerate scenarios in which we argue that budding speciation could influence our conclusions about the tempo and mode of trait evolution when using PCMs.

### 1) When evolutionary changes are concentrated at or near lineage origination

The central distinction between budding and bifurcation is the age contrast between progenitor and daughter lineage immediately after divergence. The daughter species will usually have a smaller population size and geographic distribution (Foote et al., 2007; Liow and Stenseth, 2007) and might undergo quick phenotypic change as they move towards a new adaptive peak (Eldredge and Gould, 1972; Gould and Eldredge, 1993; Hunt et al., 2008; De Lisle et al., 2021). In contrast, progenitor lineages might show a slowdown in trait evolution due to prolonged time under a stable adaptive zone (Goldberg and Foo, 2020; De Lisle et al., 2021). If lineage age is related to the tempo of trait evolution, such that younger lineages are expected to show faster rates of trait change, we would expect relatively more evolution to happen in a daughter lineage when compared to its progenitor. Thus, the disparity between two descendants of a budding node in a phylogenetic tree should not be attributed to equal amounts of change at each branch because budding suggests evolution will be concentrated in the daughter lineage (Fig. 1 top left panel).

**Figure 1:**
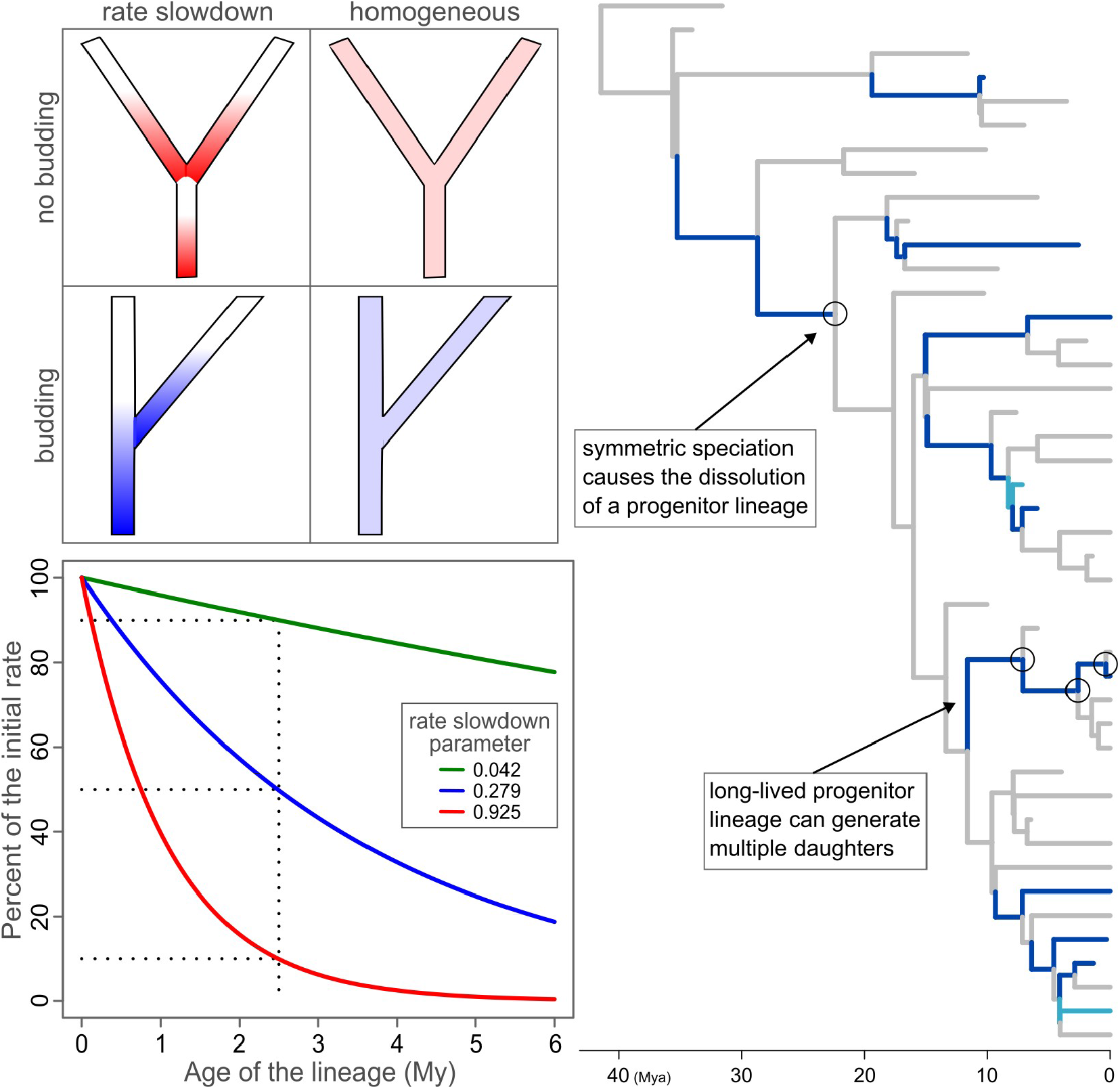
Approach used to illustrate and simulate budding speciation and trait evolution under lineage-age dependent rates. Top-left: Interaction between age-dependent trait evolution and speciation with and without budding. Stronger shades of color indicate faster rates of trait change. Bottom-left: Rate slowdown as a function of lineage age under the three treatments used. Right: Example of budding history with 20 extant species showing progenitor lineages in light and dark blue.

### 2) When daughter lineage survival depends on being ecologically different from its progenitor

The asymmetry in age generated by budding speciation can influence the competitive strength of daughter lineages relative to their progenitors and, as a result, also the extinction risk of the younger lineage (Ezard et al., 2011; Rosenblum et al., 2012; Carrillo et al., 2020; Januario and Quental, 2021). Progenitor lineages are expected to have larger population sizes and geographic ranges (Anacker and Strauss, 2014) which, everything else being equal, improves their chance of survival in interspecific competition with newly formed species. When competition between progenitor and daughter lineages is present, daughter lineages that have lived enough to be sampled, either in the fossil record or still living today, are expected to be sufficiently distinct from their progenitors to have escaped competitive exclusion (De Lisle et al., 2021). Of course, competition is not exclusive to budding. However, budding could potentially intensify the effect of interspecific competition, and eventually increase heterogeneity in trait evolution.

Although we predict an intensifying effect of budding speciation on interspecific competition, other natures of interactions might be more complex. Nuismer and Harmon (2015) demonstrated mathematically the effect of the mode of trait evolution and phylogenetic diversity (PD) in the outcome of interspecific interactions in communities of closely related taxa. They showed that PD is a good predictor of interspecific interactions if these are dependent on phenotypic matching, such as competition, with more closely related lineages showing stronger interspecific interactions. Budding could change the relationship between PD and expected trait similarity, because long-lived progenitor species would accumulate fewer evolutionary changes than expected under a homogeneous trait evolution model, such as Brownian motion, causing the role of the phylogeny as a predictor to become less prevalent. In contrast, Nuismer and Harmon (2015) show that under stabilizing coevolution the phylogeny is a poor predictor of interactions, and we do not expect that budding would influence this result.

## How do budding speciation and lineage-age-dependent processes influence estimates of trait evolution?

We use simple simulations to illustrate different scenarios in which budding speciation should impact trait evolution and, more importantly, discuss if these deviations hinder our understanding of phenotypic evolution using phylogenetic trees. We explored the impact of budding on the parameter estimates and adequacy of PCMs for continuous and discrete traits. We also investigated how likely is budding speciation to produce erroneous estimates of ancestral states. We focused our attention on phylogenies of extant species, which are most often estimated using molecular data, and in the absence of fossil tips.

### Simulation of trait evolution under budding

We simulated 50 phylogenetic trees using a constant rate birth-death model (λ = 0.2, μ = 0.1) with root age set to 40 My and excluding all extinct lineages. To reduce variation in tree size we used rejection sampling to keep only phylogenies with 250 to 350 extant lineages. We used the same pool of 50 trees to perform all simulation replicates and conducted pairwise comparisons across scenarios. We simulated a single continuous trait using a Brownian motion model (σ^2^ = 0.2) and a discrete trait with three states using an equal rates Markov model (transition rates = 0.02). To emulate a scenario of fast evolution in younger lineages we introduced a rate slowdown process. Rates of change, for both continuous and discrete traits, varied along the branches of the tree following a scaling factor (*s*) computed as a function of lineage-age (*a*), such that

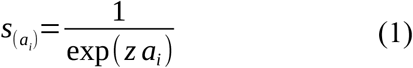

where *a*_*i*_is the average lineage-age at time interval *i* and *z* is the parameter controlling the rate slowdown. In order to compute *a*, we divided the branches of the tree into *i* time intervals of length 1x10^−3^ of the tree height. At lineage-age of 0 My, for instance, *s* is equal to 1 and it decays as a function of *z* (Fig. 1). We simulated three scenarios of lineage-age dependent rates of trait evolution: a mild effect (*z* = 0.042); a medium effect (*z* = 0.279); and a strong effect (*z* = 0.925). The parameter values were chosen to produce a rate reduction of 10%, 50%, and 90% of the base rate when a lineage becomes 2.5 My old, respectively. Because the base rate is scaled by *s*, which depends on the lineage age and the budding history of each phylogeny, the average rate of trait evolution can vary. We standardized the average rate across the tree (for both discrete and continuous traits) to differentiate the rate slowdown within lineages from the confounding effect of an overall change in the average rate. For that, we computed the weighted average as

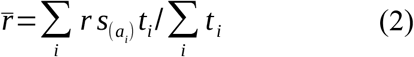

where *r* is the base rate of trait evolution (i.e., the σ^2^ for the BM model and the transition rate for the equal rates Markov model), and 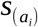 is the slowdown scale factor at a time interval *t*_*i*_ (see Equation (1)). Then we chose *r* values that minimized the distance of 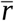 among replicates.

We simulated budding speciation using an independent binary variable to control the presence of budding on each node of the tree. As a result, long-lived progenitor lineages are produced by random events of successive budding events. We produced four scenarios of budding speciation, with frequencies of 0%, 25%, 50%, and 100% of the nodes. We also explored the effect of cladogenetic changes on discrete traits (associated or not with budding). Cladogenetic changes were simulated as a change with equal probability to any state immediately after speciation. When budding is present, cladogenetic changes were restricted to daughter lineages whereas it could happen to any lineage in the absence of budding. We also explored a scenario in which cladogenetic changes restricted to daughter lineages (thus dependent on budding) produce convergence among all daughters of the same long-lived progenitor lineage (see examples in Figs. 2 and 3). A detailed report of the simulation is available in the Supplementary Materials.

**Figure 2:**
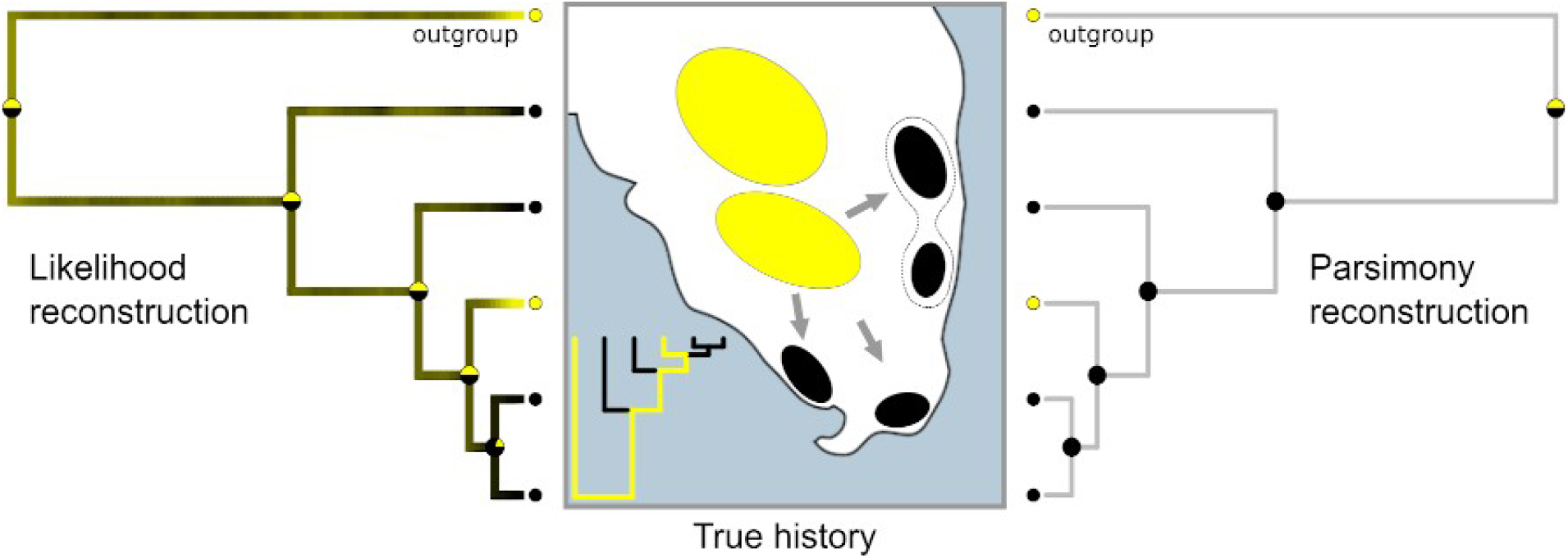
Conflict between ancestral state estimation and the true history of lineage evolution under budding. New species appeared through budding and converged three times into a different niche, associated with the coastal distribution (black). The progenitor lineage is the sister to the outgroup and carries the inland distribution (yellow). Neither likelihood nor parsimony recovered the true history of the lineages. Left: Marginal ancestral state estimation for the best fit Markov model. Branches are painted following a posterior distribution of 100 stochastic maps. Right: Most parsimonious ancestral estimation following Sankoff’s (1975) algorithm. Center: True history of the trait. Phylogeny in the bottom left shows the continuation of the progenitor lineage through multiple budding events.

**Figure 3:**
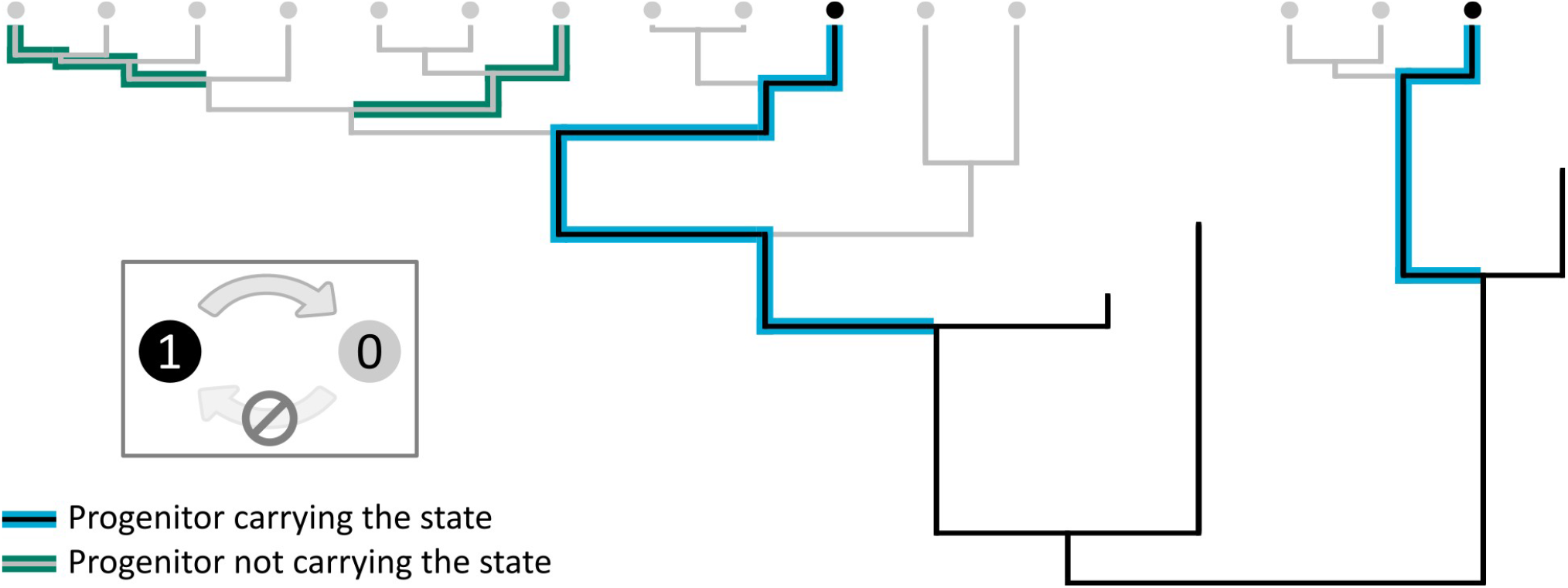
Evolution of a truly irreversible trait. Long-lived progenitor lineages carry the ancestral state whereas daughter lineages have lost the trait independently several times. Fossil lineages show evidence of trait homology. PCMs in the absence of the fossil record would wrongly estimate two independent origins of the “black” trait.

### Evaluating model adequacy and errors in ancestral estimation

We used the method described by Pennell and colleagues (2015) to evaluate the adequacy of PCMs for continuous traits in the presence of budding speciation and lineage-age-dependent rates of trait evolution. This method computes a pool of summary statistics (see Table 1 in Pennell *et al*., 2015) and compares each with an expected distribution estimated from the data. If the model is adequate, the observed summary statistics should fall around the mean of the null distribution whereas values outside the 95% highest density interval indicate that the PCM is inadequate. To evaluate the effect of budding in the ancestral estimation of discrete states we used an index of how incorrect the estimate at a node is with respect to the true history of the trait. We measured the highest marginal probability among all states excluding the true state for the nodes as an estimate of “wrongness”. This metric reflects how likely the state of a node would be estimated as the wrong ancestral state. Note that this is distinct from uncertainty because wrongness is maximized when we have certainty of the ancestral state but it is incongruent with the true (simulated) history. Uncertainty is a lesser problem than wrongness because we will not, or at least should not, support or refute evolutionary hypotheses based on uncertain estimates. Wrongness, on the other hand, can result in misleading interpretations. We then used linear mixed models to test for the association between node age and wrongness across all simulation scenarios and selected the best model using the Akaike Information Criterion (AIC).

### Effects of budding and lineage-age dependent processes on the adequacy of continuous trait evolution models

After simulating continuous traits under 12 scenarios, varying the strength of the lineage-age dependent process and the frequency of budding speciation, we estimated parameters for single rate Brownian motion (Felsenstein, 1973), variable rate BM (Eastman et al., 2011), single optimum Ornstein-Uhlenbeck (Butler and King, 2004), and Early-Burst (Harmon et al., 2010) models as implemented in *geiger* (Pennell et al., 2014). Note that none of those models is the true model that generated the data. Our goal is to evaluate which is the preferred model among the suite of PCMs most used in the literature and to better understand the potential effects of budding speciation on our inferences. We also hope that this simple illustration through the use of simulations motivates further research in model development. Overall, variable rate BM models showed significant improvement in model adequacy under budding.

Model adequacy tests for a homogeneous rate BM model (Pennell et al., 2015) detected a negative slope of the linear fit between node depth and the size of the phylogenetic independent contrasts (S_hgt_) indicating that larger trait changes are estimated to have happened closer to the tips (Fig. S3). However, a variable rate BM model does not show evidence for such deviation (Fig. S3), suggesting this is an effect of underestimating the rate variation introduced by budding (which introduces heterogeneity in a different way than the variable rate BM models). A regression of phylogenetic independent contrasts (PICs) and their expected variance (S_var_) shows that nodes connected by short branches are associated with more trait change (Fig. S2), independently if the fitted BM model was homogeneous or not. Inadequacy of S_var_ is expected due to the concentration of rates early in the history of lineages and the stronger effect of slowdown on the more longevous lineages when compared with short-lived ones. A pattern that is expected under budding speciation and punctuated equilibrium. The deviances for S_hgt_ and S_var_ are only detectable when the rate slowdown is strong, meaning that a relaxed rates model (Eastman et al., 2011) seems to be able to adequately describe trait variation if lineage-age effects are mild, but not if they are strong. Inadequacies in both S_hgt_ and S_var_ point to trait changes concentrated close to the tips, which is expected since molecular trees have an accumulation of nodes near the present, some of these generated by budding, producing new lineages with higher rates of trait evolution. Model inadequacy could be an artifact of unobserved speciation events deeper in the tree but results remain constant when we replicate analyses including extinct lineages. Deviations of S_hgt_, S_var_, and C_var_ (coefficient of variation of PICs, a measure of rate heterogeneity) get stronger as the intensity of the lineage-age-dependent slowdown factor increases. On the other hand, changing the frequency of budding speciation, while controlling for the strength of the slowdown factor, did not change the patterns of model adequacy across all summary statistics we investigated (Figs. S2-4). We did not verify any deviation from the remaining summary statistics adopted by Pennell and colleagues (2015). In summary, if budding speciation is frequent and there is strong age-dependent trait evolution (punctuated equilibrium representing an extreme version of this), current inference methods would have trouble adequately capturing patterns of trait evolution.

With respect to the support for alternative trait evolution models (i.e., Brownian motion, Ornstein-Uhlenbeck, and Early-Burst) as a function of lineage-age dependent rate variation, the OU model shows a marked increase in AIC weights in response to stronger slowdown factors (Figs. S5 and S6). In the majority of cases, the phylogenetic half-life was estimated to be multiple times longer than the age of the clade (40 My) indicating processes indistinguishable from Brownian motion (Cooper et al. 2016). Average phylogenetic half-life estimated across replicates was only shorter than clade age when the strongest slowdown factor was applied (Fig. S7). This means that an OU process is only supported when progenitor lineages are practically in stasis whereas virtually all trait evolution is concentrated on the origination of new lineages following budding speciation. In other words, if budding speciation produces a pattern congruent with Punctuated Equilibrium we expect support for OU models. These results remain constant regardless of the frequency of budding speciation used to simulate the data (Fig. S4) or the inclusion of extinct lineages.

When we simulate continuous traits under a lineage-age dependent process, the BM model with varying rates adequately describes most characteristics of the data but fails to capture the concentration of trait changes on shorter branches (S_var_). If lineage-age dependent processes happen in nature, our results reinforce the cautionary note that parameter estimates can be more informative than model choice alone (Cooper et al. 2016). Although the majority of scenarios supported OU models, only the strongest case of rate slowdown resulted in patterns distinguishable from Brownian motion. Fortunately, the deviance of the slope of absolute contrasts as a function of their expected variance (S_var_ - Pennel et al. 2015) can help to detect a concentration of trait change towards shorter branches, even when controlled for rate variation, which is one of the expectations of lineage-age-dependent rates of evolution. Those simulations are far from being comprehensive (our goal was not to be exhaustive but to provide examples to support our narrative), but they emphasize the potential effects of not explicitly considering budding speciation, in particular when age-dependent trait evolution is present. Future simulation studies should more deeply focus on the different aspects superficially touched here as well as on others not discussed.

### Effects of budding speciation and lineage-age dependent processes on ancestral estimation

Here we investigate how budding affects our estimations of ancestral state for discrete traits. As expected, all fitted models show a strong positive association between node age and wrongness (Fig. 4), meaning that ancestral estimation of nodes closer to the root of the tree is more likely to be misleading. The best-ranked linear mixed model using AIC (Table S1) suggests that budding speciation has a significant effect on wrongness when cladogenetic trait changes are also present (see example in Fig. 3). Without cladogenetic changes, there is no detectable difference between the null model (homogeneous rates and bifurcating speciation) and the model with budding speciation (Fig. 4). Budding associated with cladogenetic changes increases the chance of misleading ancestral state estimation, especially for younger nodes. This result is somewhat unexpected and important because younger nodes are often expected to have more information than older ones (Schultz et al., 1996; Boyko and Beaulieu, 2021).

**Figure 4:**
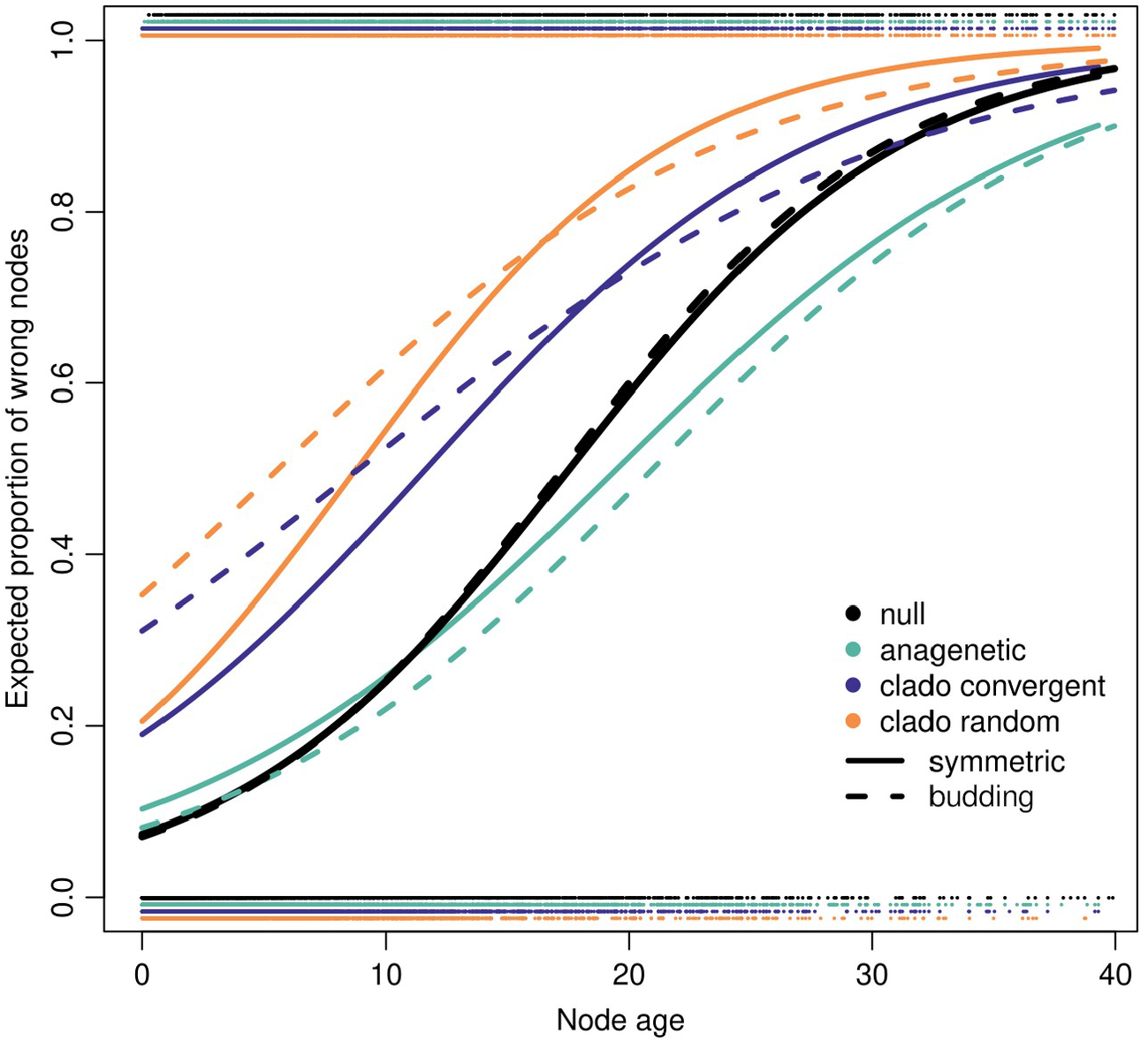
Relationship between the presence of nodes with wrongly reconstructed states and node age. Nodes were considered wrongly reconstructed if the marginal ancestral state probability for any state distinct from the correct state were > 0.5 at that node. Logistic regressions were performed with the pooled results from the 50 simulation replicates per group and independent for each type of node and study group (see **Table S1**). Dashed lines represent nodes with budding speciation whereas solid lines show nodes without budding. Dots on the top and bottom represent pooled nodes across simulation replicates correctly and incorrectly estimated, respectively.

## Can budding help us understand complex histories of trait evolution?

Here we used simulations to exemplify the effect of budding speciation in PCMs of trait evolution. Our initial results show that budding has an impact but does not completely hinder the utility of the most popular models of trait evolution. Some of the effects we report here, for the most part, can be translated as heterogeneity in trait evolution among lineages. Different from other sources of heterogeneity (e.g., Uyeda and Harmon, 2014; Boyko and Beaulieu, 2021), budding produces variation dependent on lineage identity, requiring the identification of progenitor lineages potentially comprising multiple contiguous branches of a phylogeny. From this perspective, we are optimistic about incorporating budding speciation into PCMs, and we hope our initial discussion on the subject motivates further research on how we can properly incorporate budding speciation and age-dependent trait evolution in PCMs. In fact, it is plausible that a portion of the intrinsic heterogeneity captured by rate-varying models, such as hidden rates models for discrete traits (Beaulieu and O’Meara, 2016; Caetano et al., 2018; Boyko and Beaulieu, 2021) and those applying reversible jump MCMC for continuous traits (Eastman et al., 2011; Rabosky et al., 2014; Uyeda and Harmon, 2014), is due to the effect of budding speciation.

Distinct from scenarios in which some predictor trait is responsible for rate shifts, budding is expected to affect trait evolution dependent on the mode of speciation. This introduces a complication because we need to reconstruct the budding history of lineages, which does not easily leave a trace on molecular phylogenies (e.g., it needs samples of multiple populations of recently diverged lineages). One potential solution is to use data augmentation (e.g., Quintero and Landis, 2020) to co-estimate budding history and trait evolution model parameters using simulations. This approach could be challenging because both the frequency of budding speciation and the location of the progenitor lineages would need to be sampled. However, our inability to pinpoint the location of progenitor lineages should not be used as an argument for ignoring its effect on trait evolution. Today we have several PCMs that are able to recover the signal of shifts in the tempo and mode of trait evolution without a priori hypotheses (Eastman, 2011; Rabosky et al., 2014; Uyeda and Harmon, 2014; Pagel et al., 2022) and, more importantly, there is evidence that such methods improve model adequacy (Rabosky and Goldberg, 2015; Beaulieu and O’Meara, 2016; Caetano et al., 2018). We suggest that budding speciation should be considered as a confounding factor akin to rate heterogeneity, which needs to be taken into account when estimating the history of trait evolution using molecular phylogenies—even if budding is not the focus of the study.

Another challenge is that progenitor lineages can produce scenarios incongruent with the most parsimonious history for a trait (e.g. Figs. 2 and 3). For example, ancestral estimates of the scenario shown in Figure 3 in the absence of fossil information would suggest, with confidence, that the trait is a convergence. This is a scenario in which PCM estimates can conflict with external evidence of homology. For instance, Pyron (2015) discusses the inference of multiple transitions from viviparity back to oviparity in snakes, based on PCMs, despite the external evidence based on development and physiology against so (Griffith et al., 2015). Pyron (2015) suggests that comparative approaches should not ignore external evidence but also that findings from phylogenetic inferences should be further investigated integratively. However, an unlikely ancestral reconstruction of parity might simply mean that the model is inadequate. For instance, budding speciation could help explain oviparous lineages nested deep into viviparous clades as descendants of long-lived progenitor lineages (see discussion in Pyron 2015). The budding speciation scenario would require many additional evolutionary transitions, but it would support the extensive knowledge about genetics, development, and physiology of snakes (see discussion in Griffith et al., 2015). In our view, when there is a clash between model estimates based on projections into millions of years in the past and biological knowledge, it is wise to review our models and ponder which important processes the model might be failing to capture, including the possibility of budding speciation.

## Closing remarks

Budding speciation might indirectly or directly impact both estimates of lineage diversification and trait evolution under PCMs widely used in the literature. Our results suggest that ignoring budding speciation when age-dependent trait evolution operates might lead to incorrect inferences such as inferring the wrong ancestral state for younger nodes. We also suggest that it might be possible, although challenging, to incorporate budding speciation into PCMs for both discrete and continuous traits. The introduction of budding speciation in comparative approaches, however, depends on the departure from the parsimony paradigm which we suspect is a barrier to the development of macroevolutionary models that can fully integrate external biological information about trait evolution. When we intuitively imagine a parsimonious trait history, we are doing so independently of what is known about the evolutionary history of the system. Reflecting on the role of comparative approaches and recognizing their limitations, especially when testing scenarios of complex trait evolution, is key to the development of alternative models that help the study of macroevolution to become a more integrative endeavor. The incorporation of budding speciation is one example of the direction we can take in improving comparative studies, and we hope our discussion motivates researchers to explore further some of these possibilities.

## References

Anacker, B. L., and S. Y. Strauss. 2014. The geography and ecology of plant speciation: range overlap and niche divergence in sister species. Proceedings of the Royal Society B: Biological Sciences 281:20132980.

Aze, T., T. H. G. Ezard, A. Purvis, H. K. Coxall, D. R. M. Stewart, B. S. Wade, and P. N. Pearson. 2011. A phylogeny of Cenozoic macroperforate planktonic foraminifera from fossil data. Biological Reviews 86:900–927.

Baker, J., A. Meade, M. Pagel, and C. Venditti. 2015. Adaptive evolution toward larger size in mammals. Proceedings of the National Academy of Sciences 112(16):5093–5098.

Baldwin, B. G. 2005. Origin of the serpentine-endemic herb Layia discoidea from the widespread L. glandulosa (Compositae). Evolution 59:2473.

Bapst, D. W. 2013. When can clades be potentially resolved with morphology? PLOS ONE 8:e62312.

Bapst, D. W., and M. J. Hopkins. 2017. Comparing cal3 and other a posteriori time-scaling approaches in a case study with the pterocephaliid trilobites. Paleobiology 43:49–67.

Bartoszek, K. 2014. Quantifying the effects of anagenetic and cladogenetic evolution. Mathematical Biosciences 254:42–57.

Bartoszek, K. 2020. A central limit theorem for punctuated equilibrium. Stochastic Models 36:473–517.

Bastide, P., C. Solís-Lemus, R. Kriebel, K. W. Sparks, and C. Ané. 2018. Phylogenetic comparative methods on phylogenetic networks with reticulations. Systematic Biology 67:800–820.

Baum, D. A., S. D. Smith, and S. S. S. Donovan. 2005. The Tree-Thinking Challenge. Science 310:979–980.

Beaulieu, J. M., and B. C. O’Meara. 2016. Detecting Hidden Diversification Shifts in Models of Trait-Dependent Speciation and Extinction. Systematic Biology 65:583–601.

Behm, J. E., Ives, A. R., and J. W. Boughman. 2010. Breakdown in postmating isolation and the collapse of a species pair through hybridization. The American Naturalist 175:11–26.

Benton, M. J., and P. N. Pearson. 2001. Speciation in the fossil record. Trends in Ecology & Evolution 16:405–411.

Bokma, F. 2002. Detection of punctuated equilibrium from molecular phylogenies. Journal of Evolutionary Biology 15:1048–1056.

Bokma, F. 2008. Detection of “Punctuated Equilibrium” by Bayesian estimation of speciation and extinction rates, ancestral character states, and rates of anagenetic and cladogenetic evolution on a molecular phylogeny. Evolution 62:2718–2726.

Bokma, F. 2010. Time, species, and separating their effects on trait variance in clades. Systematic Biology 59:602–607.

Boyko, J. D., and J. M. Beaulieu. 2021. Generalized hidden Markov models for phylogenetic comparative datasets. Methods in Ecology and Evolution 12:468–478.

Butler, M. A., and A. A. King. 2004. Phylogenetic comparative analysis: a modeling approach for adaptive evolution. American Naturalist 164:683–695.

Caetano, D. S., B. C. O’Meara, and J. M. Beaulieu. 2018. Hidden state models improve state-dependent diversification approaches, including biogeographical models. Evolution 72:2308–2324.

Carrillo, J. D., S. Faurby, D. Silvestro, A. Zizka, C. Jaramillo, C. D. Bacon, and A. Antonelli. 2020. Disproportionate extinction of South American mammals drove the asymmetry of the Great American Biotic Interchange. Proceedings of the National Academy of Sciences 117:26281–26287.

Clavel, J., and H. Morlon. 2017. Accelerated body size evolution during cold climatic periods in the Cenozoic. Proceedings of the National Academy of Sciences 114:4183–4188.

Cooper, N., G. H. Thomas, C. Venditti, A. Meade, and R. P. Freckleton. 2016. A cautionary note on the use of Ornstein Uhlenbeck models in macroevolutionary studies. Biological Journal of the Linnean Society 118:64–77.

Crawford, D. J. 2010. Progenitor-derivative species pairs and plant speciation. Taxon 59:1413–1423.

Crouch, N. M. A., S. M. Edie, K. S. Collins, R. Bieler, and D. Jablonski. 2021. Calibrating phylogenies assuming bifurcation or budding alters inferred macroevolutionary dynamics in a densely sampled phylogeny of bivalve families. Proceedings of the Royal Society B: Biological Sciences 288:20212178.

De Lisle, S. P., D. Punzalan, N. Rollinson, and L. Rowe. 2021. Extinction and the temporal distribution of macroevolutionary bursts. Journal of Evolutionary Biology 34:380–390.

Duchen, P., M. L. Alfaro, J. Rolland, N. Salamin, and D. Silvestro. 2021. On the effect of asymmetrical trait inheritance on models of trait evolution. Systematic Biology 70:376–388.

Eastman, J. M., M. E. Alfaro, P. Joyce, A. L. Hipp, and L. J. Harmon. 2011. A novel comparative method for identifying shifts in the rate of character evolution on trees. Evolution 65:3578–3589.

Eldredge, N., and S. J. Gould. 1972. Punctuated equilibria: An alternative to phyletic gradualism, pp. 82–115. In: Schopf, T. J. M., ed. Models in Paleobiology. Freeman, Cooper and Co.; San Francisco, California.

Elliot, M. G., and A. Ø. Mooers. 2014. Inferring ancestral states without assuming neutrality or gradualism using a stable model of continuous character evolution. BMC Evolutionary Biology 14.

Ezard, T. H. G., T. Aze, P. N. Pearson, and A. Purvis. 2011. Interplay between changing climate and species’ ecology drives macroevolutionary dynamics. Science 332:349–351.

Felsenstein, J. 1973. Maximum likelihood estimation of evolutionary trees from continuous characters. American Journal of Human Genetics 25:471–492.

Felsenstein, J. 1981. Evolutionary trees from DNA sequences: A maximum likelihood approach. Journal of Molecular Evolution 17:368–376.

Felsenstein, J. 1985. Phylogenies and the Comparative Method. American Naturalist 125:1–15.

Foote, M. 1996. On the Probability of ancestors in the fossil record. Paleobiology 22:141–151.

Foote, M., J. S. Crampton, A. G. Beu, B. A. Marshall, R. A. Cooper, P. A. Maxwell, and I. Matcham. 2007. Rise and Fall of Species Occupancy in Cenozoic Fossil Mollusks. Science 318:1131–1134.

Funk, D. J., and K. E. Omland. 2003. Species-Level Paraphyly and Polyphyly: Frequency, Causes, and Consequences, with Insights from Animal Mitochondrial DNA. Annual Review of Ecology, Evolution, and Systematics 34:397–423.

Goldberg, E. E., and J. Foo. 2020. Memory in Trait Macroevolution. The American Naturalist 195:300–314.

Gottlieb, L. D. 2004. Rethinking classic examples of recent speciation in plants. New Phytologist 161:71–82.

Gould, S. J., and N. Eldredge. 1993. Punctuated equilibrium comes of age. Nature 366:223–227.

Grant, V. 1981. Plant speciation, 2nd ed. New York, NY, USA: Columbia University Press.

Griffith, O. W., D. G. Blackburn, M. C. Brandley, J. U. Van Dyke, C. M. Whittington, and M. B. Thompson. 2015. Ancestral state reconstructions require biological evidence to test evolutionary hypotheses: A case study examining the evolution of reproductive mode in squamate reptiles. Journal of Experimental Zoology Part B: Molecular and Developmental Evolution 324:493–503.

Hagen, O., K. Hartmann, M. Steel, and T. Stadler. 2015. Age-dependent speciation can explain the shape of empirical phylogenies. Systematic Biology 64:432–440.

Harmon, L. J., J. B. Losos, T. J. Davies, R. G. Gillespie, J. L. Gittleman, W. B. Jennings, K. H. Kozak, M. A. McPeek, F. Moreno-Roark, T. J. Near, A. Purvis, R. E. Ricklefs, D. Schluter, J. A. Schulte II, O. Seehausen, B. L. Sidlauskas, O. Torres-Carvajal, T. J. Weir, and A. Ø. Mooers. 2010. Early bursts of body size and shape evolution are rare in comparative data. Evolution 64: 2385–2396.

Helmstetter, A.J., Zenil-Ferguson, R., Sauquet, H., Otto, S.P., Méndez, M., Vallejo-Marin, M. et al. (2023) Trait-dependent diversification in angiosperms: Patterns, models and data. Ecology Letters 26:640–657.

Huelsenbeck, J. P., R. Nielsen, and J. P. Bollback. 2003. Stochastic mapping of morphological characters. Systematic Biology 52:131–158.

Hunt, G., M. A. Bell, and M. P. Travis. 2008. Evolution toward a new adaptive optimum: phenotypic evolution in a fossil stickleback lineage. Evolution 62:700–710.

Janzen, T., F. Bokma, and R. S. Etienne. 2022. Nucleotide substitutions during speciation may explain substitution rate variation. Systematic biology 71:1244–1254

Januario, M., and T. B. Quental. 2021. Re-evaluation of the “law of constant extinction” for ruminants at different taxonomical scales. Evolution 75:656–671.

Lackey, A. C, J. W. Boughman. 2017. Evolution of reproductive isolation in stickleback fish. Evolution 71:357–372

Liow L.H. & Stenseth N.C. 2007 The rise and fall of species: implications for macroevolutionary and macroecological studies. Proceedings of the Royal Society of London, Series B 274:2745–2752.

Mallet, J., N. Besansky, and M. W. Hahn. 2016. How reticulated are species? BioEssays 38:140–149.

Mattila, T. M., and F. Bokma. 2008. Extant mammal body masses suggest punctuated equilibrium. Proceedings of the Royal Society B: Biological Sciences 275:2195–2199.

May, M. R., and B. R. Moore. 2020. A bayesian approach for inferring the impact of a discrete character on rates of continuous-character evolution in the presence of background-rate variation. Systematic Biology 69:530–544.

Mayr, E. 1942. Systematics and the origin of species. New York, NY, USA: Columbia University Press.

Meier, R., and R. Willmann. 2000. The Hennigian species concept. In Wheeler, Q. D., and R. Meier (eds). Species concepts and phylogenetic theory: a debate (pp. 30–43). Columbia University Press, New York.

Monroe, M. J., and F. Bokma. 2009. Do speciation rates drive rates of body size evolution in mammals? The American Naturalist 174:912–918.

Nuismer, S. L., and L. J. Harmon. 2015. Predicting rates of interspecific interaction from phylogenetic trees. Ecology Letters 18:17–27.

O’Meara, B. C. 2012. Evolutionary Inferences from Phylogenies: A Review of Methods. Annual Review of Ecology, Evolution, and Systematics 43:267–285.

Otero, A., P. Vargas, M. Fernández-Mazuecos, P. Jiménez-Mejías, V. Valcárcel, I. Villa-Machío, and A. L. Hipp. 2022. A snapshot of progenitor–derivative speciation in Iberodes (Boraginaceae). Molecular Ecology 31:3192–3209.

Pagel, M., C. O’Donovan, and A. Meade. 2022. General statistical model shows that macroevolutionary patterns and processes are consistent with Darwinian gradualism. Nature Communications 13:1113.

Papuga, G., P. Gauthier, V. Pons, E. Farris, and J. D. Thompson. 2018. Ecological niche differentiation in peripheral populations: a comparative analysis of eleven Mediterranean plant species. Ecography 41:1650–1664.

Parins-Fukuchi, C. 2021. Morphological and phylogeographic evidence for budding speciation: an example in hominins. Biology Letters 17:20200754.

Patsis, A., R. P. Overson, K. A. Skogen, N. J. Wickett, M. G. Johnson, W. L. Wagner, R. A. Raguso, J. B. Fant, and R. A. Levin. 2021. Elucidating the evolutionary history of oenothera sect. Pachylophus (onagraceae): A phylogenomic approach. Systematic Botany 46:799–811.

Pennell, M. W., and L. J. Harmon. 2013. An integrative view of phylogenetic comparative methods: Connections to population genetics, community ecology, and paleobiology. Annals of the New York Academy of Sciences 1289:90–105.

Pennell, M. W., J. M. Eastman, G. J. Slater, J. W. Brown, J. C. Uyeda, R. G. FitzJohn, M. E. Alfaro, and L. J. Harmon. 2014. geiger v2.0: An expanded suite of methods for fitting macroevolutionary models to phylogenetic trees. Bioinformatics 30:2216–2218.

Pennell, M. W., R. G. FitzJohn, W. K. Cornwell, and L. J. Harmon. 2015. Model adequacy and the macroevolution of angiosperm functional traits. American Naturalist 186:E33–E50.

Philippe, H., and C. J. Douady. 2003. Horizontal gene transfer and phylogenetics. Current Opinion in Microbiology 6:498–505.

Pyron, R. A. 2015. Advancing perspectives on parity-mode evolution. Journal of Experimental Zoology Part B: Molecular and Developmental Evolution 324:562–563.

Quintero, I., and M. J. Landis. 2020. Interdependent phenotypic and biogeographic evolution driven by biotic interactionst. Systematic Biology 69:739–755.

Rabosky, D. L., S. C. Donnellan, M. Grundler, and I. J. Lovette. 2014. Analysis and visualization of complex macroevolutionary dynamics: An example from australian scincid lizards. Systematic Biology 63:610–627.

Rabosky, D. L., and E. E. Goldberg. 2015. Model inadequacy and mistaken inferences of trait-dependent speciation. Systematic Biology 64(2):340–355.

Raup, D. M. 1985. Mathematical models of cladogenesis. Paleobiology 11:42–52.

Raup, D. M., and S. J. Gould. 1974. Stochastic simulation and evolution of morphology-towards a nomothetic paleontology. Systematic Zoology 23(3):305–322.

Richmond, J. Q. and E. L. Jockusch. 2007. Body size evolution simultaneously creates and collapses species boundaries in a clade of scincid lizards. Proceedings of Biological Sciences 274:1701–1708.

Rosenblum, E. B., B. A. J. Sarver, J. W. Brown, S. Des Roches, K. M. Hardwick, T. D. Hether, J. M. Eastman, M. W. Pennell, and L. J. Harmon. 2012. Goldilocks meets Santa Rosalia: An ephemeral speciation model explains patterns of diversification across time scales. Evolutionary Biology 39:255–261.

Sankoff, D. 1975. Minimal mutation trees of sequences. SIAM Journal on Applied Mathematics 28:35–42.

Schultz T.R., Cocroft R.B., Churchill G.A. 1996. The reconstruction of ancestral character states. Evolution 50:504–511.

Taylor, E. B., J. W. Boughman, M. Groenenboom, M. Sniatynski, D. Schluter, and J. L. Gow. 2006. Speciation in reverse: Morphological and genetic evidence of the collapse of a threespined stickleback (Gasterosteus aculeatus) species pair. Molecular Ecology 15:343–355.

Uyeda, J. C., and L. J. Harmon. 2014. A novel Bayesian method for inferring and interpreting the dynamics of adaptive landscapes from phylogenetic comparative data. Systematic Biology 63:902–918.

Wagner, P. J. 1998. A likelihood approach for evaluating estimates of phylogenetic relationships among fossil taxa. Paleobiology 24:430–449.

