## Supplementary material for "How important is budding speciation for comparative studies?"

Daniel S. Caetano[1,2] and Tiago Bosisio Quental[2]
[1] *Department of Biological Sciences, Towson University, Towson, MD, 21252, USA*
[2] *Department of Ecology, University of São Paulo, São Paulo, SP, 05508-090, Brazil*

---

### TABLE OF CONTENTS

#### List of Tables

**Table S1:** Logistic regression models for the relationship between the presence of nodes with wrongly reconstructed (discrete) states and node age.

#### List of Figures

**Figure S1:** Branching history of lineages interpreted through bifurcation or budding.
**Figure S2:** Effect of lineage-age dependent trait evolution and frequency of budding speciation on expected variance ($S_{var}$).
**Figure S3:** Effect of lineage-age dependent trait evolution and frequency of budding speciation on node depth ($S_{hgt}$).
**Figure S4:** Effect of lineage-age dependent trait evolution and frequency of budding speciation on absolute contrasts ($C_{var}$).
**Figure S5:** Support for alternative trait evolution models for scenarios with 25% of budding speciation events and in the presence of lineage-age dependent rates with varying decaying factors.
**Figure S6:** Support for alternative trait evolution models for the null scenario without the effect of lineage-age-dependent rates of trait evolution.
**Figure S7:** Phylogenetic half-life as a function of budding frequency and strength of the lineage-age dependent slowdown factor.

#### List of Sections

**Section 1:** Simulating phylogenetic trees
**Section 2:** The lineage-age dependent process
**Section 3:** Simulating budding speciation
**Section 4:** Working with discrete states
*Section 4.1: Cladogenetic changes associated with budding speciation*
**Section 5:** Working with continuous states

**References**

**Table S1:** Relationship between the presence of nodes with wrongly reconstructed (discrete) states and node age. Logistic regressions were performed with the pooled results from 50 simulation replicates (see Fig 4). The first column shows the exact model configuration implemented in the R package *lme4* (Bates et al., 2015). The variable names are as follows; *wrongness* : Nodes were considered wrongly reconstructed if the marginal ancestral state probability for any state distinct from the correct state were > 0.5 at that node (nodes reconstructed with < 0.5 marginal ancestral state probability were considered uncertain). *node_age* : Age of the node, with older nodes closer to the root. *sim_type* : One of 12 distinct simulation scenarios with a combination of the frequency of budding speciation (0%, 25%, 50%, and 100%) and strength of the rate slowdown as a function of lineage-age (z = 0.042, 0.279, and 0.925). *node_type* : If budding speciation or not (frequency of budding speciation varied depending on the scenario). *descendant_history* : The type of history of the clade descendant of a budding event. We separated clades by the presence or absence of further budding events among the descendants of a budding node and if the budding event leads to a terminal branch on the phylogeny or not. The value of AIC for the best model was 30119.47. ΔAIC: difference from best model AIC. DF: degrees of freedom.

| **Linear mixed model ( as used in R )** | **Included effects** | **ΔAIC** | **DF** |
|---|---|---|---|
| glmer(wrongness ~ node_age + sim_type + (1\|node_type) + (1\|descendant_history) , family = binomial) | Simulation type, if nodes represent budding events, and the history of the descendants of budding events | 0.00 | 7 |
| glmer(wrongness ~ node_age + sim_type + (1\|node_type) , family = binomial) | Simulation type and whether the nodes represent budding or symmetric speciation | 148.02 | 6 |
| glmer(wrongness ~ node_age + sim_type + (1\|descendant_history) , family = binomial) | Simulation type and the history of clades descendant from budding events | 40.95 | 6 |
| glm(wrongness ~ node_age + sim_type , family = binomial) | Simulation type | 203.78 | 5 |
| glm(wrongness ~ node_age , family = binomial) | No effects | 2571.34 | 2 |

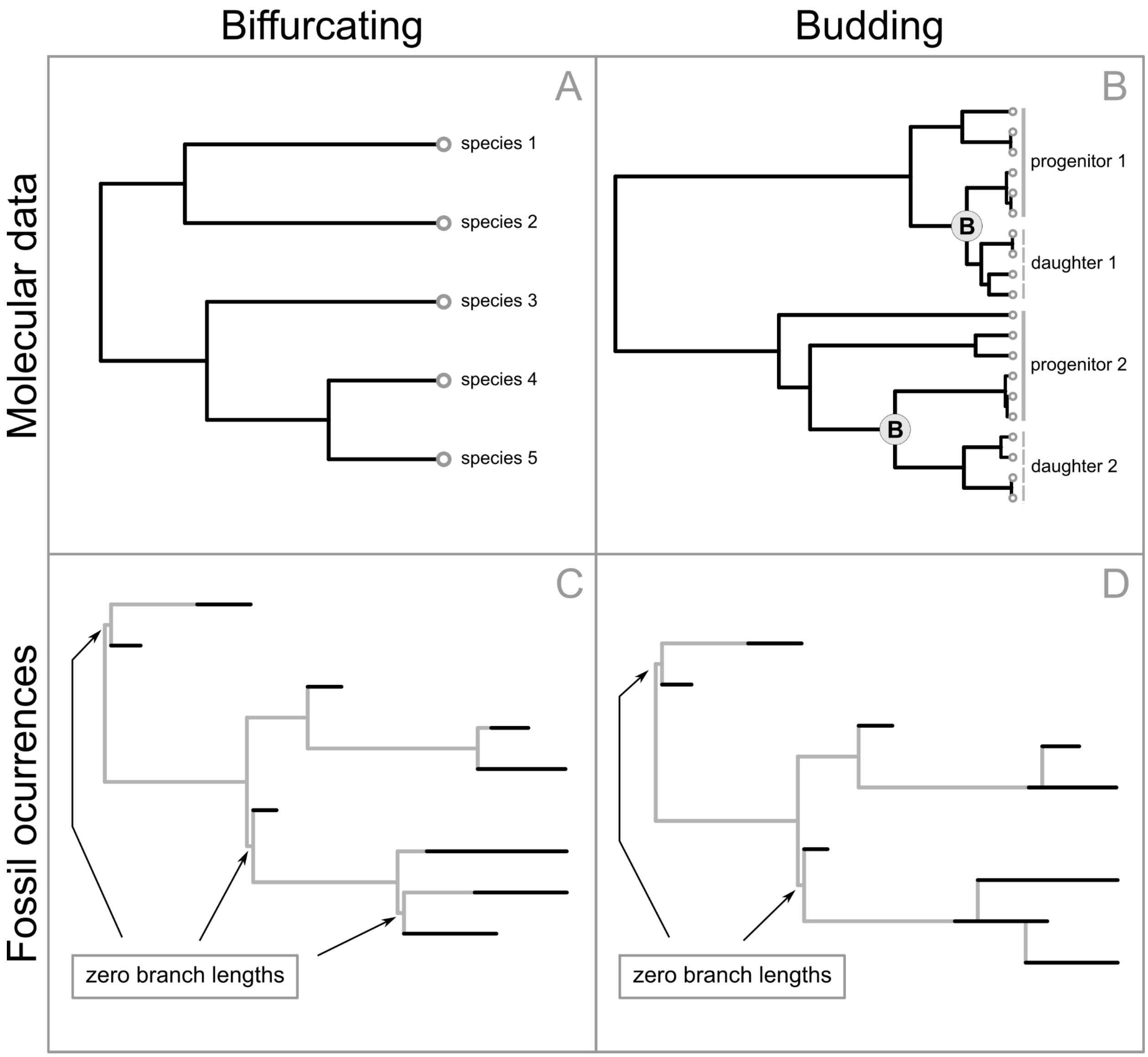


**Figure S1:** Branching history of lineages interpreted through bifurcation or budding. A) Bifurcating molecular tree shows no evidence of budding. B) Molecular tree showing budding speciation. Internal nodes labeled with "B" show recent budding events causing the progenitor species to become paraphyletic. C) Bifurcating fossil phylogeny with "ghost" lineages representing estimated longevity beyond fossil occurrences. D) Same fossil phylogeny as in C), but with budding events inferred from the fossil history. Budding decreases the total length of "ghost" branches and minimizes the number of zero branch lengths added due to incongruence between topology and age of fossil lineages.

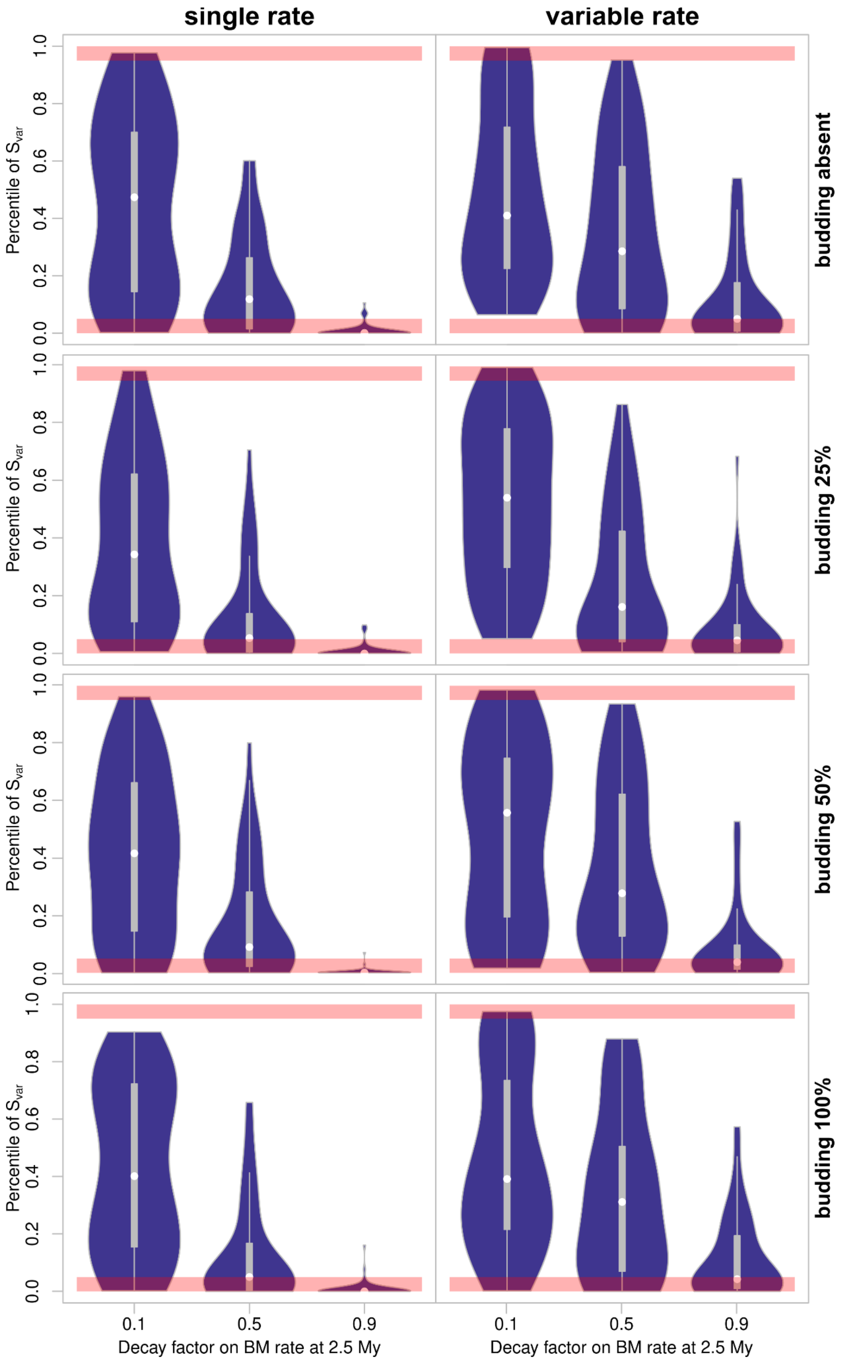


**Figure S2:** Effect of lineage-age dependent trait evolution and frequency of budding speciation on estimates of a Brownian Motion with homogeneous rate (left column) and a variable rates BM model (right column). Plots show the distribution of the percentile of $S_{var}$ across 50 simulation replicates. Each observation is the slope of absolute contrasts as a function of their expected variance ($S_{var}$). Each percentile is computed from a null distribution of $S_{var}$ generated following Pennel *et al.*, 2015. Values outside the 95% density (regions highlighted in salmon) are evidence of model inadequacy. The x axes show the slowdown factor ($\boldsymbol{\sigma}_{2.5}^2 = \boldsymbol{\sigma}_0^2 - \boldsymbol{\sigma}_0^2 * $ decay factor). Small values of $S_{var}$ indicate slowdown causes a concentration of trait changes in short branches.

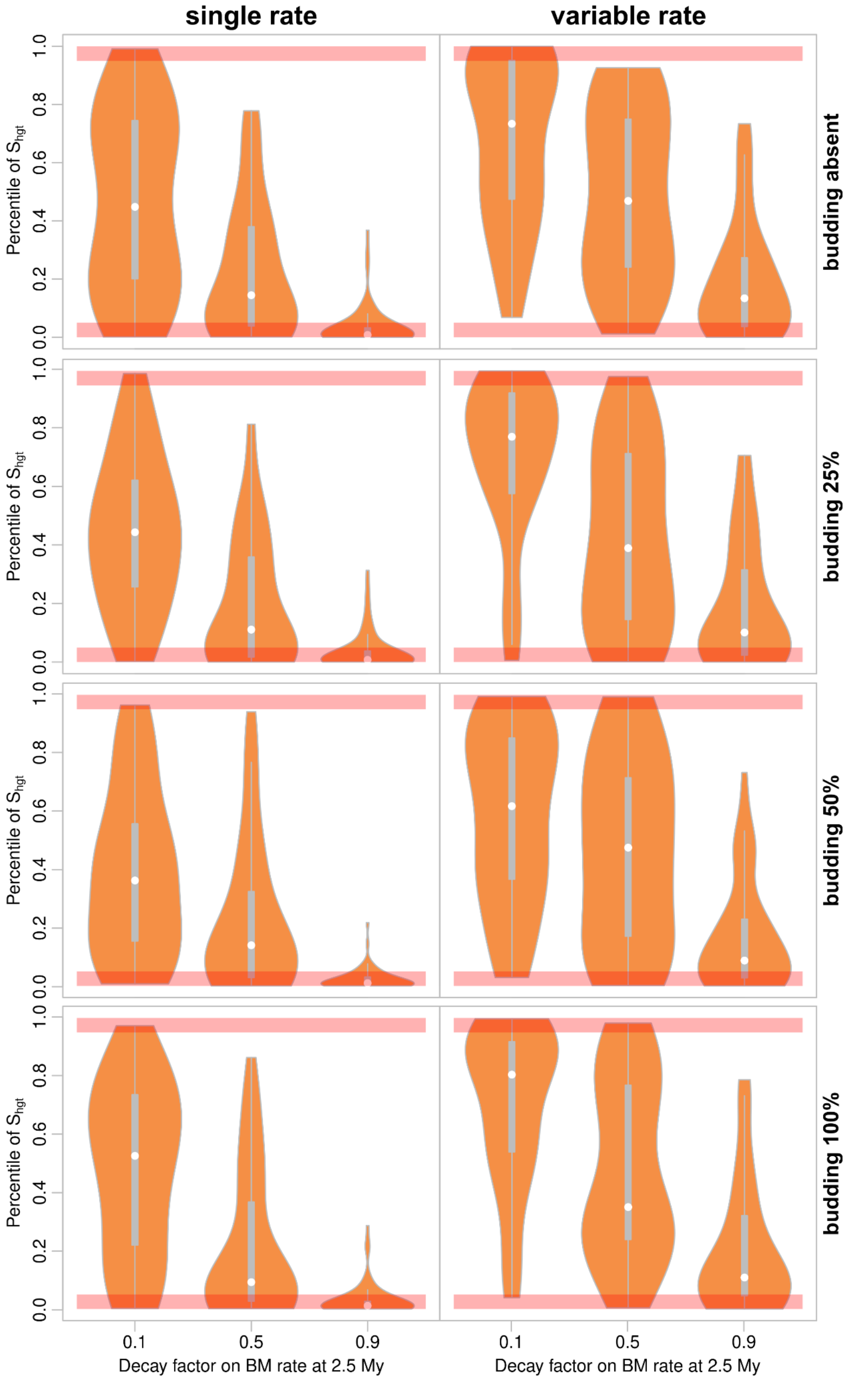


**Figure S3:** Effect of lineage-age dependent trait evolution and frequency of budding speciation on estimates of a Brownian Motion with homogeneous rate (left column) and a variable rates BM model (right column). Plots show the distribution of the percentile of $S_{hgt}$ across 50 simulation replicates. Each observation is the slope of absolute contrasts as a function of node depth ($S_{hgt}$). Each percentile is computed from a null distribution of $S_{hgt}$ generated following Pennel *et al.*, 2015. Values outside the 95% density (regions highlighted in salmon) are evidence of model inadequacy. The x axes show the slowdown factor ($\boldsymbol{\sigma}_{2.5}^2 = \boldsymbol{\sigma}_0^2 - \boldsymbol{\sigma}_0^2 *$ decay factor). Small values represent negative slopes and indicate that more trait change is estimated to have happened at younger nodes. When allowing rates to vary, this relationship fades away.

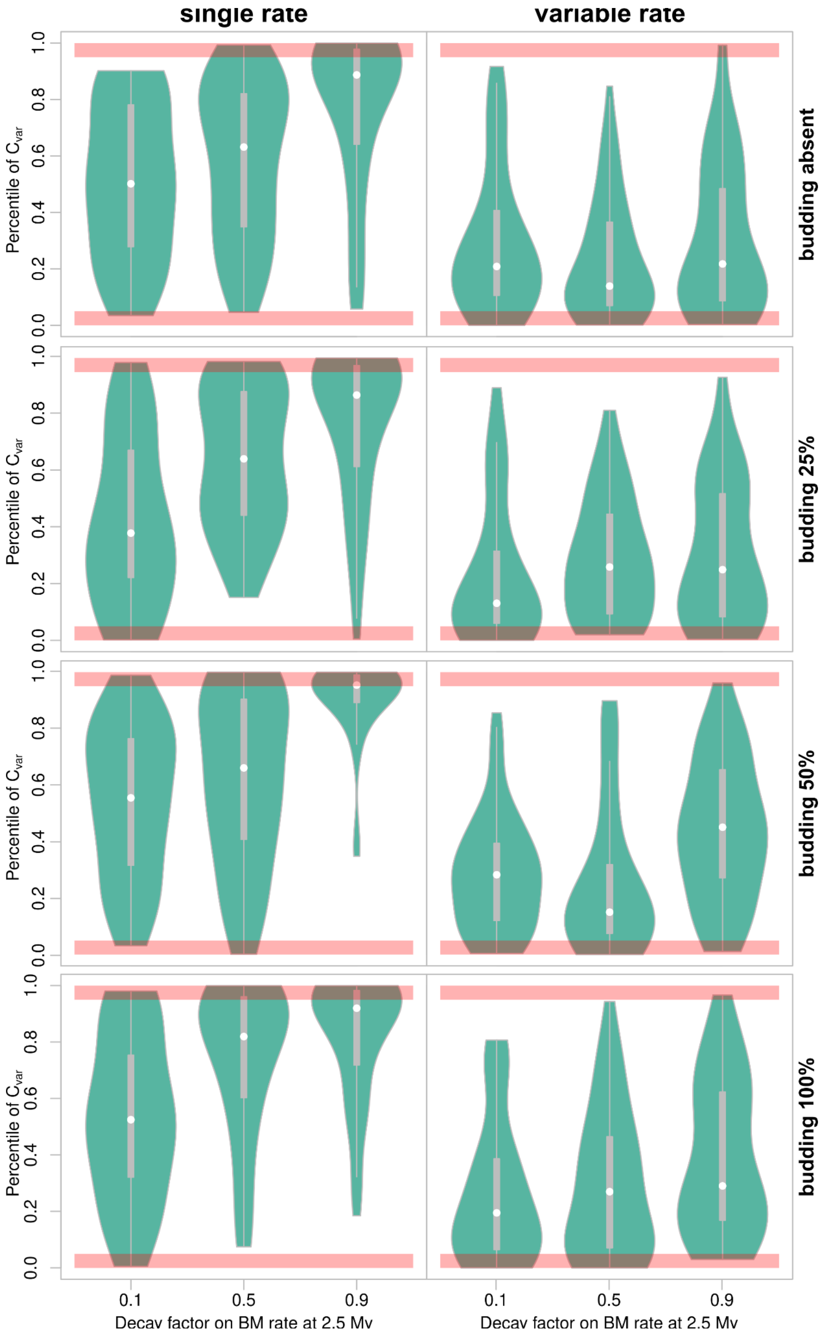


**Figure S4:** Effect of lineage-age dependent trait evolution and frequency of budding speciation on estimates of a Brownian Motion with homogeneous rate (left column) and a variable rates BM model (right column). Plots show the distribution of the percentile of $C_{var}$ across 50 simulation replicates. Each observation is the coefficient of variation of absolute contrasts ($C_{var}$). Each percentile is computed from a null distribution of $C_{var}$ generated following Pennel *et al*., 2015. Values outside the 95% density (regions highlighted in salmon) are evidence of model inadequacy. The x axes show the slowdown factor ($\boldsymbol{\sigma}_{2.5}^2 = \boldsymbol{\sigma}_0^2 - \boldsymbol{\sigma}_0^2$ * decay factor). High values indicate the model is not properly accounting for rate variation.

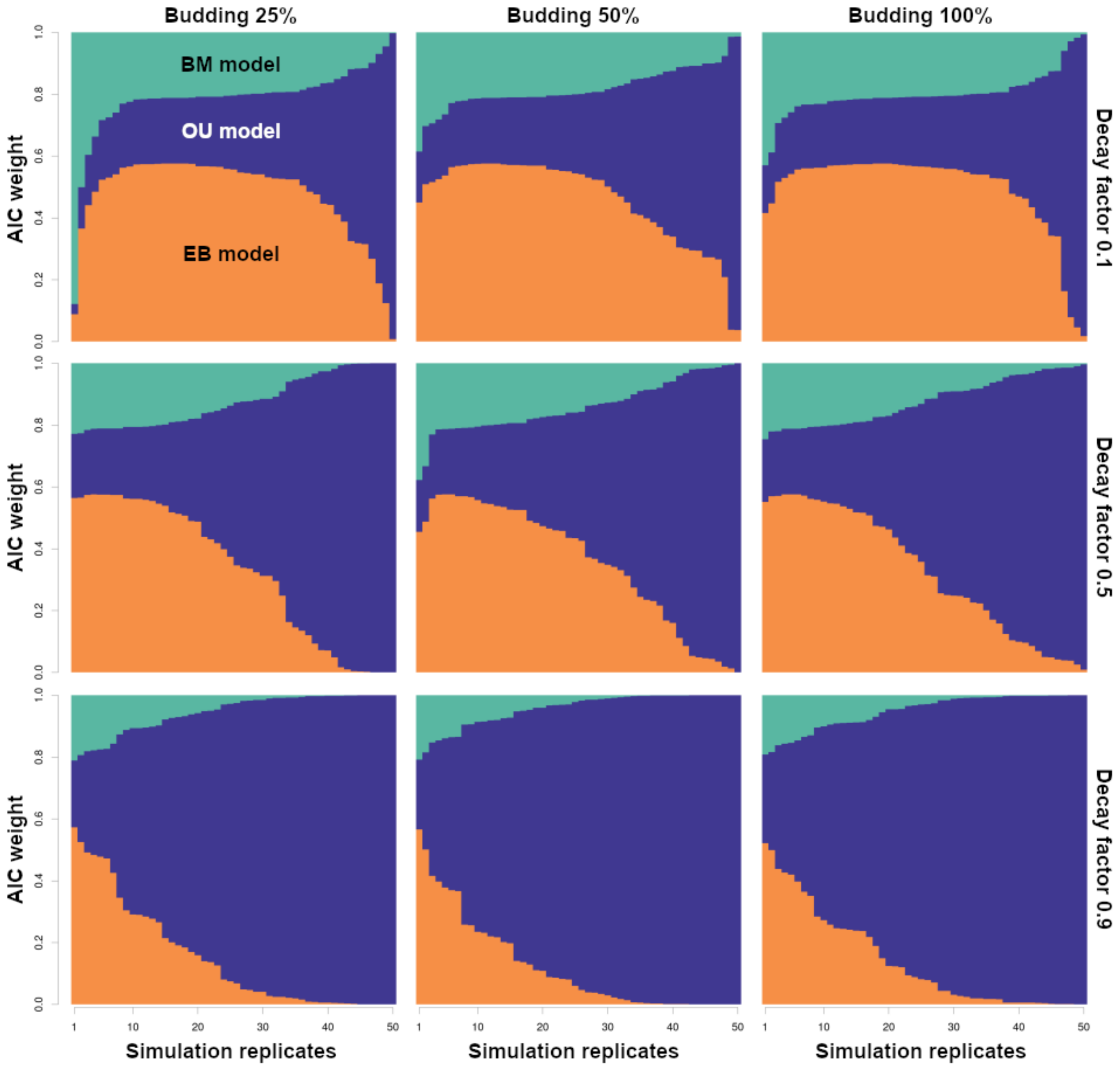


**Figure S5:** Support for alternative trait evolution models for scenarios with 25%, 50%, and 100% of budding speciation events and in the presence of lineage-age dependent rates with varying decaying factors (0.1, 0.5, and 0.9). Panels show the distribution of AIC weights for Brownian motion (BM - teal), Ornstein-Uhlenbeck (OU - blue), and Early Burst (EB - orange) models across simulation replicates. Simulation replicates were ordered within each simulation scenario by the support for each model to aid in the visualization of patterns (i.e., simulation number 1 is not the same across all panels, for example). Support for OU models increases as relative rates of trait evolution concentrate at speciation events. The overall pattern remains the same independent of the frequency of budding speciation. See Figure S6 for a scenario with no effect of lineage-age-dependent rate of trait evolution.

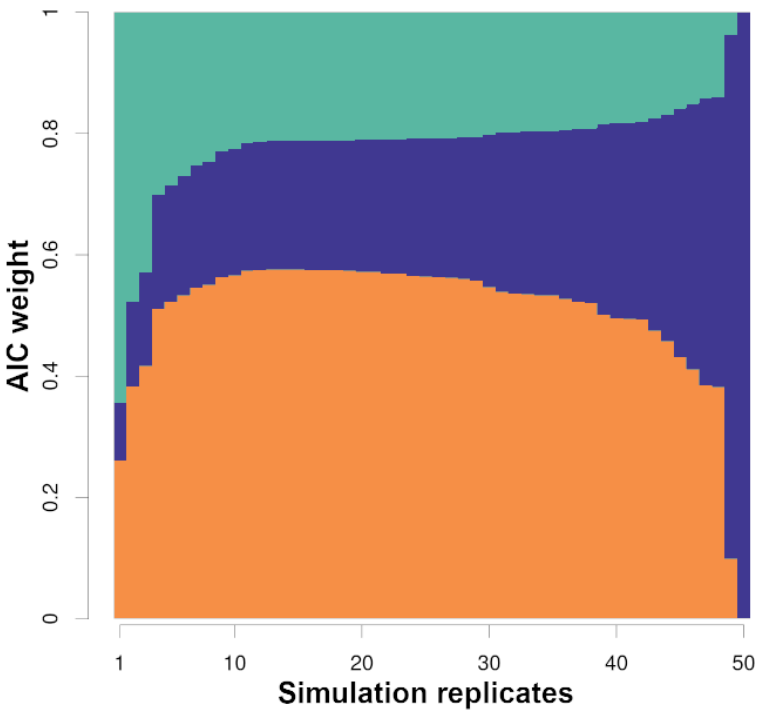


**Figure S6:** Support for alternative trait evolution models for the null scenario without the effect of lineage-age-dependent rates of trait evolution. Because rates are homogeneous, the result is the same regardless of the presence and frequency of budding speciation. The figure shows the distribution of AIC weights for Brownian motion (BM - teal), Ornstein-Uhlenbeck (OU - blue), and Early Burst (EB - orange) models across simulation replicates. Simulation replicates were ordered by the support for each model to aid in the visualization.

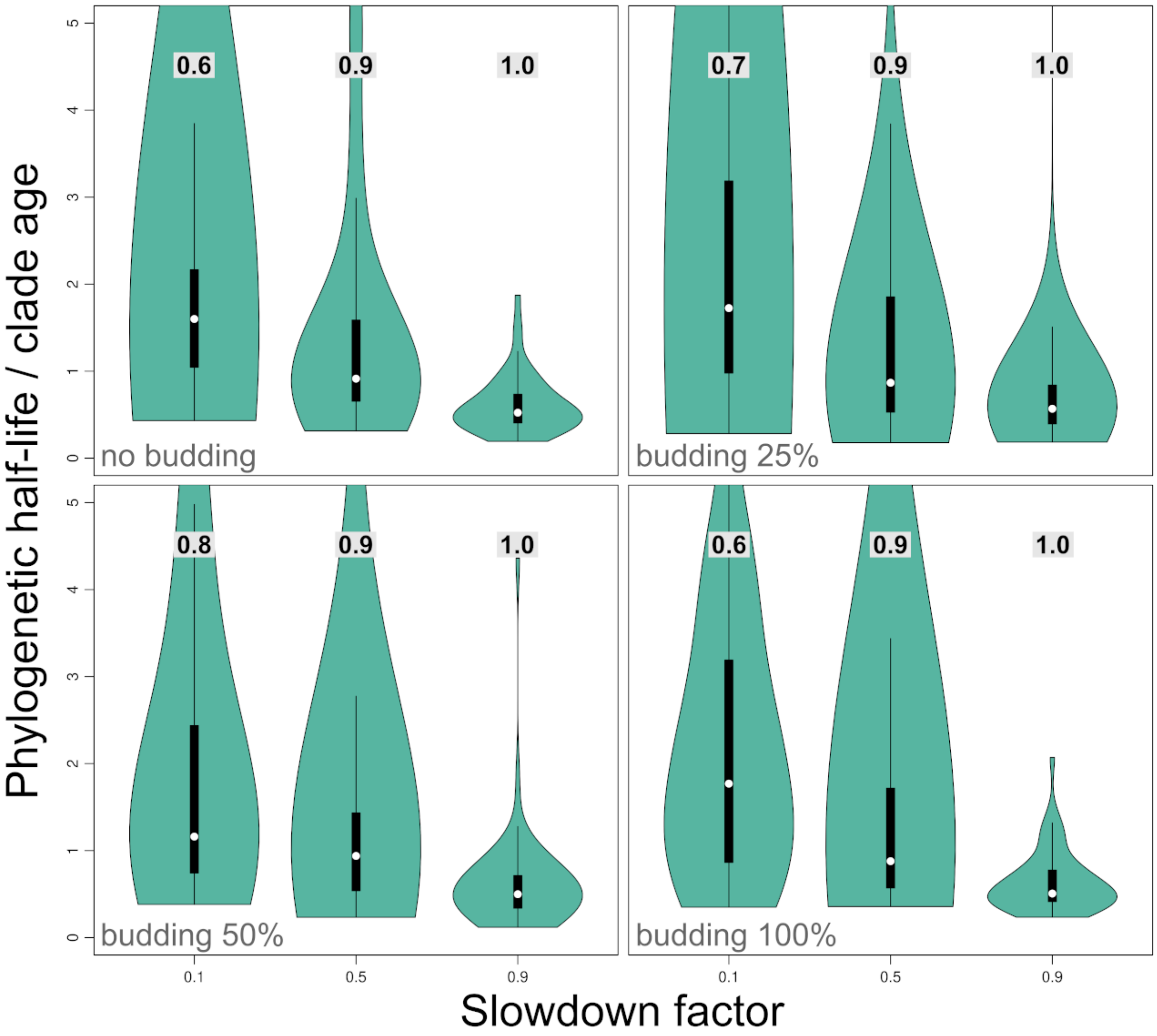


**Figure S7:** Distribution of phylogenetic half-life estimated from Ornstein-Uhlenbeck models fitted to multiple simulation replicates with increasing budding frequency and strength of the lineage-age dependent slowdown factor. Phylogenetic half-life was rescaled to represent the proportion of the clade age, thus a value of 1 indicates a half-life equal to the tree's age (40 My). Values > 1 represent a very weak pull towards the optima, with half-lives multiple times the age of the clade indicating a process indistinguishable from unbounded trait evolution. Numbers in gray boxes show the proportion of the total pool of replicates that favored OU models.

***Section 1: Simulating phylogenetic trees***

We simulated 50 phylogenetic trees using a birth-death process ($\lambda = 0.2$, $\mu = 0.1$) and fixed the root age equal to 40 Mya. We used rejection sampling to keep only trees between 250 to 350 extant lineages in order to avoid biases due to significant variations in tree size. We excluded all extinct lineages in order to emulate the case of molecular phylogenies of extant species. However, we repeated all estimates using complete phylogenies and did not find any significant bias in the results due to the inclusion of extant lineages only. We repeated each of the simulation scenarios across each of the 50 simulated trees. As a result, all comparisons were made pairwise. Simulation of the budding speciation process (see Section 3) does not change tree topology or branch lengths. Thus, the trees remained fixed.

***Section 2: The lineage-age dependent process***

The lineage-age dependent model of trait evolution simulates the scenario in which lineages are more likely to show rapid trait change earlier after speciation and then gradually reduce the pace of trait evolution as they approach a stable evolutionary optimum. Put simply, the rate of trait evolution is a function of the age of the lineage—as lineage age increases the rate of trait evolution decreases. This relationship is not restricted to linear functions; more complex functions (such as the exponential decay used in this study) can also be implemented.

We implemented a lineage-age-dependent rate of trait evolution by rescaling the branches of the tree. Faster rates of trait change are emulated by extending the branch whereas shortening branches reduce the expected amount of trait change. We rescaled branches based on the average lineage age at a very small time interval, computed with the age of the lineage at the start and end points of the time interval. Then, we modeled the slowdown in rates of trait evolution as a function of lineage age with an exponential distribution. We controlled the mean of the exponential distribution in order to create scenarios with varying levels of rate slowdown, from patterns that are indistinguishable from a homogeneous rate to a strong concentration of rates after the birth of new lineages, similar to the expectations under punctuated equilibrium. This approach allows for the use of most univariate models of trait evolution available for both discrete and continuous traits without any modification to existing software packages.

***Section 3: Simulating budding speciation***

We simulate budding speciation by assigning a binary variable to each node of the phylogeny. A value equal to 1 represents a divergence through budding (asymmetric) speciation and 0 indicates a symmetric (bifurcation) event. We used a binary distribution to control the frequency of budding speciation. The probability of budding is independent for each node of the tree, so long-lived progenitor lineages are the result of random events rather than autocorrelation in the probability of budding speciation after a previous budding event happened.

Budding speciation does not affect the simulation of lineages under a birth-death model. Rates of birth and death remain the same regardless of the mode of speciation (if symmetric or budding). As a result, here we simulate phylogenetic trees independent of budding and create budding scenarios a posteriori by assigning each node of the tree as a budding event or not. Because budding histories are created using fixed trees, here we make all 50 simulation replicates across the scenarios pairwise, meaning that a single tree can be used to create budding scenarios with varying characteristics. The advantage of this approach is that we can compare scenarios of budding speciation and trait evolution while keeping the phylogeny constant.

When a budding speciation event happens, one of the descendant lineages is a daughter lineage and the other is the progenitor lineage. We randomly selected which of the lineages is the progenitor (either the branch on the left or on the right of a node). The progenitor lineage is the continuation of the ancestral lineage which survives the speciation event and inherits the age of the ancestor whereas the daughter lineage represents a newly formed species with an age of 0 My. The progenitor lineage can generate another daughter lineage if the descendant node is also defined as a budding speciation event by the simulation. When that happens, then the progenitor lineage branch is the continuation of the ancestral branch, creating a long-lived continuous progenitor lineage that can span multiple nodes of the tree.

***Section 4: Working with discrete states***

For discrete traits, we used a symmetric rate Makov transition matrix for three states

$$Q = \begin{matrix} -0.06 & 0.03 & 0.03 \\ 0.03 & -0.06 & 0.03 \\ 0.03 & 0.03 & -0.06 \end{matrix}$$

. We randomly assigned the state at the root of the tree. Preliminary tests with binary states suggested that the rapid decay of information for ancestral state reconstruction of older nodes would significantly constrain the window of parameter values that we could explore using simulations. Thus we chose to use three states instead of two.

*Section 4.1: Cladogenetic changes associated with budding speciation*

We implemented cladogenetic models of trait evolution for discrete traits associated with budding speciation. Cladogenetic change emulates the scenario in which a new species is formed through budding and undergoes rapid trait change whereas the progenitor species remain relatively unchanged. Rapid trait change of newly formed lineages can be a reflection of competition because only lineages that are distinct enough in key traits to explore a new adaptive optimum might survive enough time to be sampled in the fossil record or be still alive today. We modeled this scenario by assigning a binary variable to each budding node to control whether there is a cladogenetic change associated with the birth of the new species.

We performed all simulations using 50% of cladogenetic change probability at every budding node of the tree. We represented a cladogenetic change as a random draw from the potential states, excluding the current trait at that point of the simulation. A random draw is adequate in this case because all discrete trait simulations used symmetric transition matrices (Q), but it is possible to weigh transitions based on asymmetric transition matrices under more complex scenarios. The new lineage was allowed to change immediately after the cladogenetic event.

***Section 5: Working with continuous states***

Simulation of continuous traits was similar to discrete traits. We simulated a single continuous trait following a Brownian Motion model with a base rate ($\sigma^2$) of 0.2 and a decrease in rate controlled by the lineage-age dependent process.

***References cited in this supplement:***

Eastman, J. M., M. E. Alfaro, P. Joyce, A. L. Hipp, and L. J. Harmon. 2011. A Novel Comparative Method for Identifying Shifts in the Rate of Character Evolution on Trees. *Evolution* 65:3578–3589.

Pennell, M. W., R. G. FitzJohn, W. K. Cornwell, and L. J. Harmon. 2015. Model adequacy and the macroevolution of angiosperm functional Traits. *American Naturalist* 186:E33–E50.
